# Plasma Oxylipin Profiling by High Resolution Mass Spectrometry Reveal Signatures of Inflammation and Hypermetabolism in Amyotrophic Lateral Sclerosis

**DOI:** 10.1101/2023.07.07.547101

**Authors:** Adriano B. Chaves-Filho, Larissa S. Diniz, Rosangela S. Santos, Rodrigo S. Lima, Hector Oreliana, Isabella F.D. Pinto, Lucas S. Dantas, Alex Inague, Rodrigo L. Faria, Marisa H.G. Medeiros, Isaías Glezer, William T. Festuccia, Marcos Y. Yoshinaga, Sayuri Miyamoto

**Author notes:** Corresponding authors: Adriano B. Chaves-Filho, Ph.D.; Sayuri Miyamoto, Ph.D.

## Abstract

Amyotrophic lateral sclerosis (ALS) is a neurodegenerative disease characterized not only by progressive loss of motor neurons, but also linked to systemic hypermetabolism, oxidative stress, and inflammation. In this context, oxylipins have been investigated as signaling molecules linked to neurodegeneration. However, the nature and role of major oxylipins involved in ALS disease progression remain unclear. Importantly, most methods focused on oxylipin analysis are based on low resolution mass spectrometry (LRMS), which usually confers high sensitivity, but not great accuracy for lipid identification, as provided by high-resolution MS (HRMS). Here, we established an ultra-high performance liquid chromatography coupled HRMS (LC-HRMS) method for simultaneous analysis of 126 oxylipins in plasma, including lipid hydroxides, ketones, epoxides, prostaglandins, leukotrienes, and others in a 15-minute run. Intra- and inter-day method validation showed high sensitivity (0.3 – 25 pg), accuracy and precision for more than 90 % of quality controls. This method was applied for the analysis of oxylipins in plasma of ALS rats overexpressing the mutant human Cu/Zn-superoxide dismutase gene (SOD1-G93A) at asymptomatic (ALS 70 days old) and symptomatic stages (ALS 120 days old), and their respective age-matched wild type controls (WT 70 days old and WT 120 days old). From the 56 oxylipins identified in plasma, 17 species were significantly altered. Remarkably, most of oxylipins linked to inflammation and oxidative stress derived from arachidonic acid, such as, prostaglandins, lipoxins, mono-hydroxides, and isoprostane, were increased in ALS 120d rats. In contrast, the linoleic acid diols involved in fatty acid uptake and β-oxidation, 9(10)-DiHOME and 12(13)-DiHOME, were strongly decreased in the ALS 120d. In summary, we developed and validated a high-throughput LC-HRMS method for oxylipin analysis and provided a comprehensive overview of plasma oxylipins involved in ALS disease progression. Noteworthy, the oxylipins altered in plasma of ALS 120d rats have potential to be investigated and used as biomarkers for inflammation and hypermetabolism in ALS.

**GRAPHICAL ABSTRACT:** 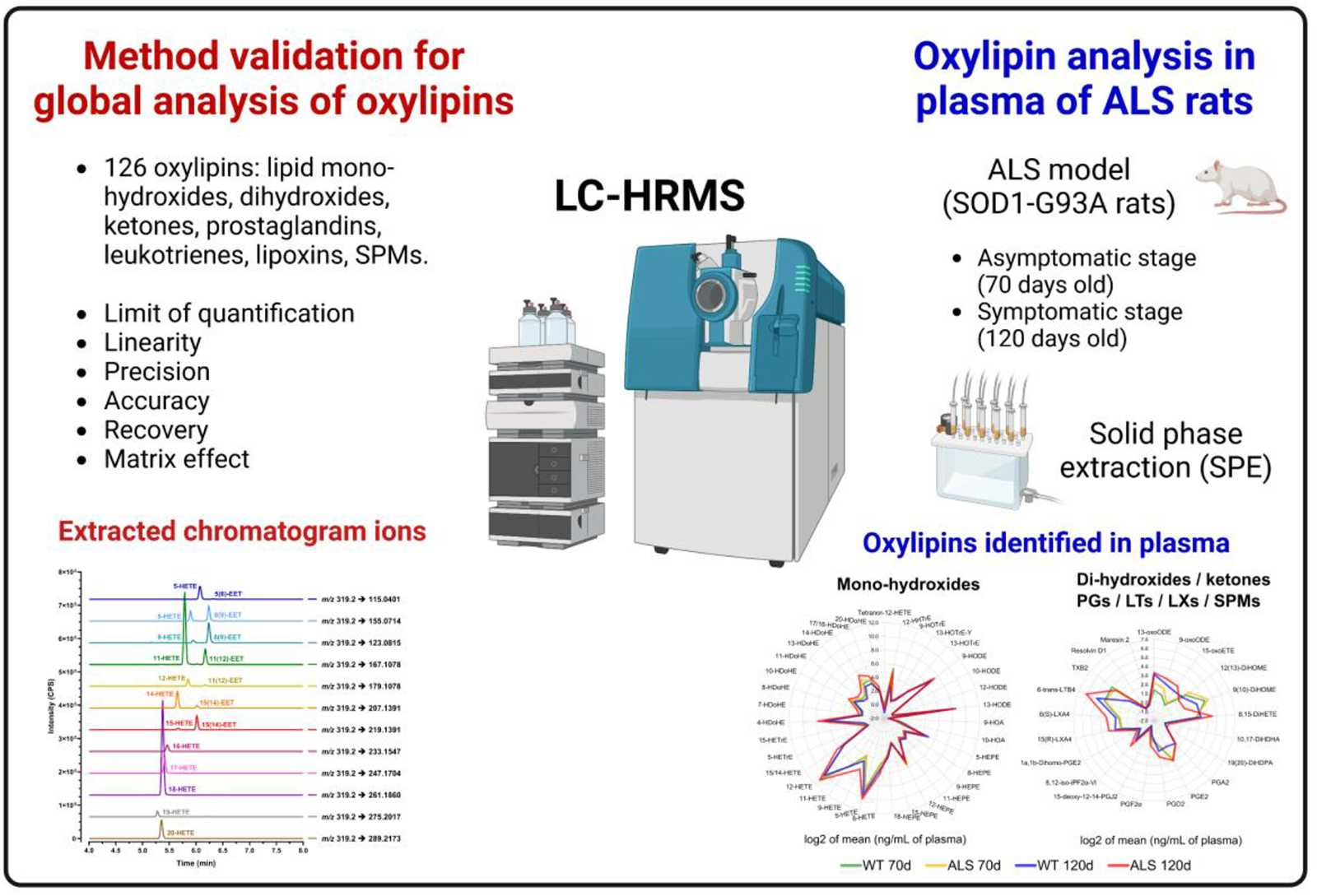

## 1. Introduction

Lipids are emerging as key regulators of fundamental cellular processes including cell survival, division, and death [1]. Among several lipid classes, oxidized lipids have gained much attention due to their association with oxidative stress, inflammation, and cell signaling [2]. Importantly, polyunsaturated fatty acids (PUFAs) are lipid targets of oxidation by enzymatic and non-enzymatic pathways. Enzymatically, PUFAs can be oxidized by lipoxygenases (LOX), cyclooxygenases (COX) and cytochrome P450 (CYP) yielding stereospecific oxylipins [3], such as leukotrienes, prostaglandins, and lipoxins [4]. On the other hand, PUFAs may be oxidized by reactive oxygen species (ROS), yielding a set of non-stereospecific products, including lipid hydroperoxides, hydroxides, ketones, aldehydes and cyclic peroxides [5,6]. According to their chemical properties, reactive electrophilic oxylipins may modify other biomolecules, thus changing their biological functions and/or propagating signals through the interaction with protein receptors [7,8]. Indeed, the biological relevance of oxylipins under pathological conditions have been extensively investigated, particularly in neurodegenerative diseases [9,10].

The amyotrophic lateral sclerosis (ALS) is a neurodegenerative disorder characterized by death of motor neurons in brain and spinal cord, which leads to muscular atrophy, paralysis, and death [11]. Importantly, alterations in lipid metabolism, chronic inflammation and oxidative stress are strongly linked to ALS disease progression [12–14]. Although it remains unclear if oxylipins play a key role in ALS, previous studies have reported changes in metabolism of some oxylipins using animal models [15,16] and samples from patients [9,17]. Not surprisingly, most of these studies have focused on the role of prostaglandin E_2_ (PGE_2_), a well-known pro-inflammatory oxylipin derived from enzymatic oxidation of arachidonic acid (AA) by COX. For example, it was reported that PGE_2_ is not only increased in cerebrospinal fluid of ALS patients [17], but it can also stimulate the release of glutamate from astrocytes, leading to increased influx of calcium and superoxide anion production in neurons [18]. Interestingly, the PGE_2_ receptor EP2 was found highly expressed in motor neurons from the spinal cord of SOD1-G93A mice [19]. Remarkably, EP2 genetic deletion improved motor strength and extended survival [19]. On the other hand, a study performed with an organotypic culture model of ALS showed that PGE_2_ paradoxically protects motor neurons from chronic toxicity induced by glutamate [20]. Mechanistically, the protective effect of the EP2 receptor was found to be dependent on the cAMP signaling [20]. Importantly, not only AA-derived prostaglandins, but also other oxylipins may play a key role in ALS. As recently reported, ALS patients showed a decrease of 13- and 9-hydroxy-octadecadienoic acids (13-HODE and 9-HODE), both oxylipins derived from linoleic acid (LA), in blood plasma as compared to control subjects [9]. However, a comprehensive and global study focused on identification of major oxylipins involved in ALS is still missing. In addition, the discovery of new oxylipin biomarkers would contribute for identification of early disease onset and monitoring of disease progression.

Advancements in mass spectrometry (MS) technologies and bioinformatic tools are enabling the identification of thousands of lipid molecular species and the characterization of their spatial and temporal changes under pathological conditions [21–23]. However, performing a global and comprehensive analysis of oxylipins is still a challenge due to their structural similarity, chemical instability, and low concentration in biological matrices (usually from pM to nM) [24]. Of note, most of the methods developed for oxylipin detection so far are based on low resolution mass spectrometry (LRMS) [25–28]. This analytical approach confers high sensitivity for detection, but not great accuracy for molecular identification of oxylipins. Indeed, the development of methods based on high resolution mass spectrometry (HRMS) could improve not only chemical characterization, but also support identification of new oxylipins. However, the validation of a comprehensive and high-throughput HRMS method for global analysis of oxylipins is still missing. Thus, efforts towards development and optimization of analytical strategies based on HRMS are essential to guarantee accurate characterization of oxylipins in biological matrices.

Here, we aimed to develop and validate an accurate and reproducible method for simultaneous analysis of 126 oxylipins based on liquid chromatograph coupled to HRMS. This analytical tool was applied in a rat model of ALS (SOD1-G93A) to access the oxylipins involved in disease progression. In summary, oxylipins altered in plasma of ALS rats at a symptomatic stage reflected oxidative stress, inflammation, and lipid hypermetabolism.

## 2. Methods

### 2.1. Materials and preparation of work solutions

Mono-hydroxides isomers derived from arachidonic, docosahexaenoic, linoleic and oleic acids were synthesized, purified, and quantified as previously described [27,29]. Other oxidized fatty acids and deuterated internal standards used for oxylipin analysis were obtained from Cayman Chemical. For analysis of reference standards, stock solutions of each analyte were prepared in methanol at a concentration of 10 µg/mL. Working solutions of all oxylipins were prepared by serial dilution of the stock solution. All solutions were stored at -20 °C before their use. Reference and internal standards used for oxylipin analysis are described in Tables 1 and 2.

**Table 1.**
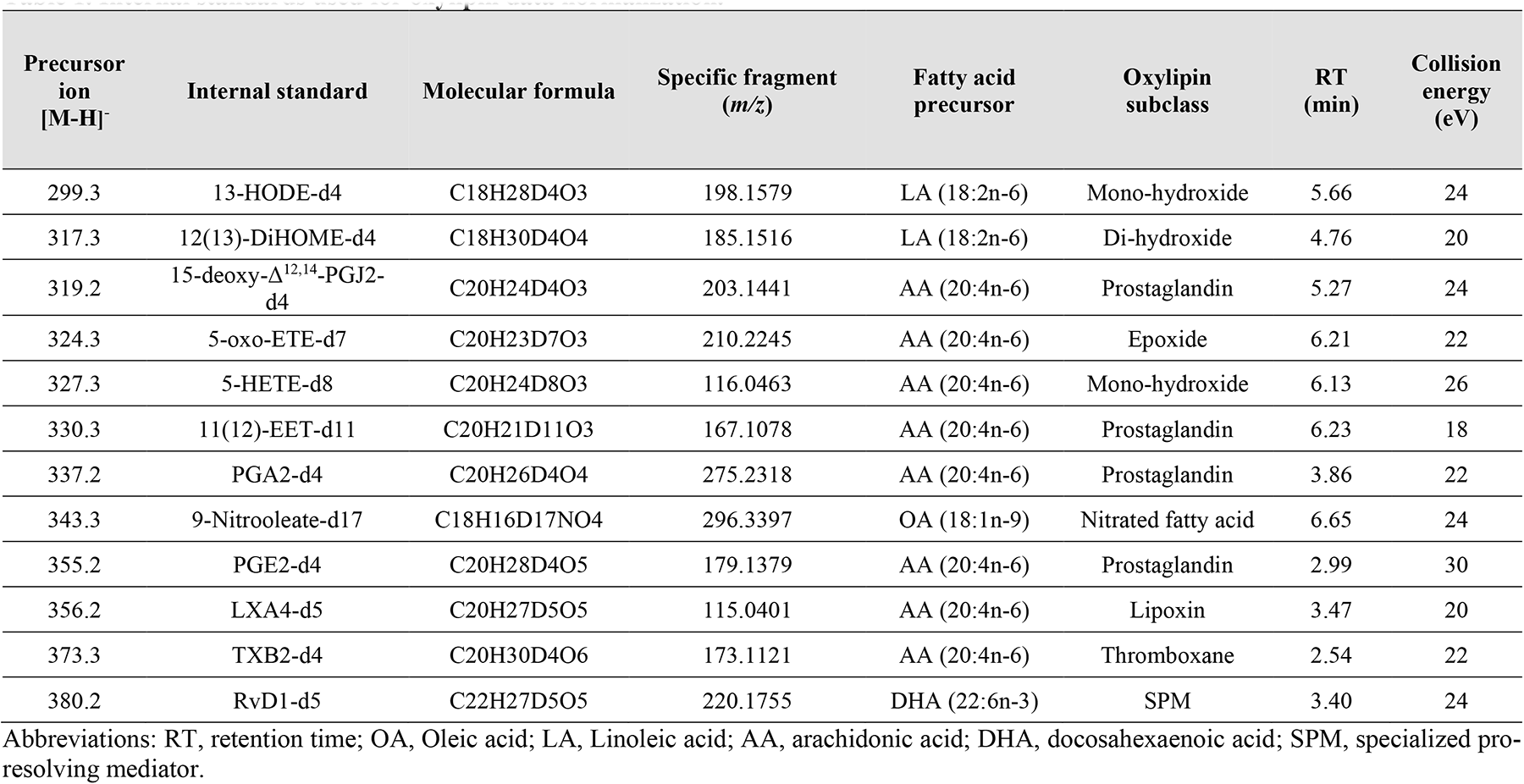
Internal standards used for oxylipin data normalization.

### 2.2. Characterization of specific fragments for quantification of oxylipins

The characterization of specific fragments of oxylipins was performed by direct infusion using a high-resolution mass spectrometry ESI-Q-TOFMS system (Triple TOF^®^ 6600, Sciex, Concord, US). Data acquisition was performed using the Analyst^®^ 1.7.1 software (Sciex). Briefly, each reference and internal standard was infused into the MS system to obtain information about the optimal collision energy (CE). The investigation of specific fragments using MS/MS data was performed with PeakView^®^ 2.2 and ChemDraw^®^ 20.0 softwares.

### 2.3. LC-MS/MS conditions for analysis of oxylipins

The chromatographic and mass spectrometry conditions for analysis of oxylipins were set as previously described [26], with modifications. Analytes were analyzed by an ESI-Q-TOFMS system (as described in the method session 2.2) interfaced with an ultra-high performance liquid chromatography (UHPLC Nexera, Shimadzu, Kyoto, Japan). The samples were loaded into a BEH^®^ (UPLC^®^ C18 column, 1.7 µm, 2.1 mm i.d. x 100 mm) with a flow rate of 0.5 mL min^-1^ and the oven temperature maintained at 35 °C. For reverse-phase LC, mobile phase A consisted of water/acetonitrile (70:30), while mobile phase B composed of acetonitrile/isopropanol (50/50). Both mobile phases A and B contained 0.01% acetic acid. Oxylipins were separated by a 15 min gradient as follows: from 0.1 to 40% B over the first 3.5 min., from 40 to 75% B from 3.5-6.0 min., from 75 to 99% B from 6.0-6.5 min., hold at 99% B from 6.5-10.5 min., decrease from 99 to 0.1% B during 10.5-11 min., and hold at 0.1% B from 11–15 min.

The MS was operated in negative ionization mode, and the scan range was set at a mass-to-charge ratio of 200-500 Da (MS) and 50-500 Da (MS/MS). Data for lipid molecular species identification and quantification was obtained by product ion targeted acquisition. Data acquisition using Analyst^®^ 1.7.1 was performed with a cycle time of 0.560 s with 100 ms acquisition time for MS scan and 10 ms acquisition time for MS/MS scan. An ion spray voltage of -4.5 kV and the cone voltage at -80 V were set to analysis. Additional parameters included curtain gas set at 25 psi, both nebulizer and heater gases at 50 psi and interface heater at 500 °C. Identification and quantification of each analyte were performed by monitoring their specific fragments using Multiquant^®^ software.

### 2.4. Linearity, accuracy, precision, and limit of quantification

Calibration curves were prepared by adding 100 µL of internal standard mixture (Table S1) to each concentration point of reference standards. The linear regressions of the area ratios (peak area of analyte/peak area of internal standard) obtained from the calibration curves were used to calculate the corresponding analyte concentration in samples. The precision and accuracy were determined for quality controls containing high, medium, and low concentration of analytes. Both precision and accuracy were checked intra-and inter-daily. The precision was expressed as a coefficient of variation (CV), calculated by dividing the standard deviation by the mean of the area ratio, and then multiplied by 100%. The accuracy was expressed by the relative error (RE), calculated by the experimental concentration obtained by applying the quality control area ratio in the curve equation. The experimental concentration was divided by the theoretical concentration, multiplied by 100%. The value obtained was subtracted by 100 to obtain the RE. The limit of quantitation (LOQ) was defined as the lower point in the curve showing signal to noise ratio greater than 7 and maximum 20% of CV and RE.

### 2.5. Recovery rate and matrix effect

The recovery rate was determined by spiking internal standards into control plasma samples and comparing the peak areas obtained after extraction with the corresponding peak areas of a reference internal standard solution. The matrix effect was carried out by spiking internal standards into plasma extracts and comparing the peak areas with the corresponding peak areas of a reference internal standard solution.

### 2.6. Animal model

Animal study was conducted in accordance with the ethical principles for animal experimentation and conducted according to the guidelines of the National Council for Animal Experimentation Control (Conselho Nacional de Controle de Experimentação Animal – CONCEA, Ministry of Science, Technology, Innovation and Communications, Brazil) under the protocol number 14/2013. Male Sprague Dawley rats overexpressing human SOD1-G93A were obtained from Taconic and bred with wild-type Sprague-Dawley females to establish a colony. Genotyping by PCR to detect exogenous hSOD1 transgene was performed by amplification of ear DNA at 20 days of age. Rats were housed under controlled laboratory conditions, including room temperature, a 12 hours light/12 hours dark cycle with food and water ad libitum. Asymptomatic ALS rats (ALS 70d; n = 6) and their wild type controls (WT 70d; n = 6) were sacrificed at 73 ± 4 days of age, while symptomatic ALS rats (ALS 120d; n = 6) and their control (WT 120d; n = 6) were sacrificed at 122 ± 6 days of age. The criterion for euthanasia of symptomatic ALS rats was loss of 15% of their maximum body weight. Rats were fasted for 6 h and anesthetized by isoflurane inhalation at a dose of 4% for induction and 2% for maintenance. Blood was collected by cardiac puncture into a tube containing heparin. Plasma was obtained after centrifugation at 2,000 x g for 10 min at 4 °C and stored at -80 °C until further processing. All animal procedures were approved by University of Sao Paulo – Chemistry Institute’s Animal Care and Use Committee under the protocol number 14/2013.

### 2.7. Analysis of oxylipins in plasma of ALS rats

Sample preparation for analysis of oxylipins was performed as previously described [26], with modifications. Briefly, 500 µL of plasma samples were added by 2 mL of cold 10 µM 3,5-di-tert-4-butylhydroxytoluene (BHT) in methanol and 100 µL of internal standards (Table S1). Samples were vortexed during 30s and maintained in ice-cold bath for 10 min. Then, samples were centrifuged at 2000 x *g* for 10 min at 4 °C. Supernatant was collected and samples were concentrated to 500 µL using N_2_ gas and maintained in ice-cold bath until purification of oxylipins by solid phase extraction (SPE). Briefly, Strata-X reversed-phase SPE columns (8B-S100-UBJ, Phenomenex) were washed with 3 mL of methanol and then equilibrated with 3 mL of ultrapure water. Then, samples were applied into SPE column. Columns were washed with 1 mL of 5% methanol and then oxylipins were eluted with 1 mL of 100% methanol. Samples were dried using N_2_ gas, dissolved in 100 µL of methanol and immediately analyzed by LC-MS/MS as described in the method session 2.3.

### 2.8. Statistical analysis

All statistical analyses were performed with Metaboanalyst (website: www.metaboanalyst.ca). In brief, data were log_10_ transformed prior to statistical analyses. All groups were compared by multivariate (principal component analysis, PCA) and univariate analyses (one-way ANOVA, followed by Tukey’s post-hoc; p < 0.05). We also constructed heatmap plots using clusters of oxylipins that were statistically altered in the different groups and clusters of samples. Graphics were generated using GraphPad Prism 9, Metaboanalyst, Excel, Chemdraw, and Inkscape softwares.

## 3. Results and discussions

### 3.1. Analytical method validation

In our study, we developed and validated a method based on liquid chromatography coupled to high-resolution mass spectrometry (LC-HRMS) for analysis of oxylipins. To our knowledge, only a few studies focused on validation of HRMS-based methods for analysis of specific oxylipins [30–32]. Of note, analytical strategies based on drift tube ion mobility coupled with HRMS were used to improve oxylipin detection [31,32]. Here, we focused on targeted analysis of 12 labeled internal standards (Table 1) and 126 oxylipins (Table 2). Of note, most of analytes monitored in our method were derived from enzymatic or non-enzymatic oxidation of arachidonic (AA), docosahexaenoic (DHA), linoleic (LA), eicosapentaenoic (EPA), and dihomo-γ-linolenic (DGLA) acids (Fig. 1A). Regarding oxylipin subclasses, we monitored lipid mono-hydroxides (41 species), prostaglandins (34 species), epoxides (13 species), specialized pro-resolving mediators (SPMs) (8 species), leukotrienes (7 species), and others (Fig. 1B).

**Figure 1.**
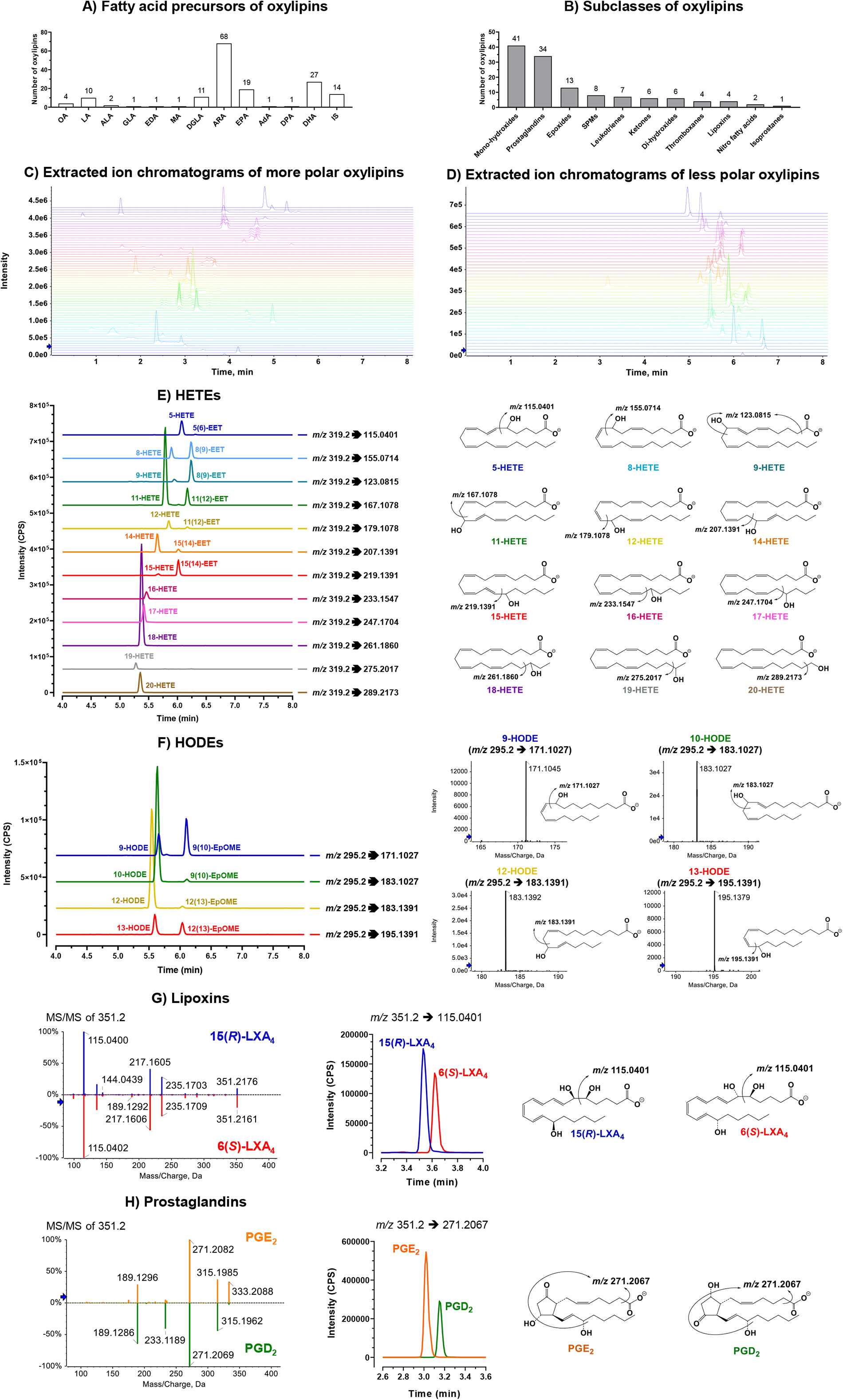
Profile of oxylipins monitored by high resolution LC-MS/MS. (A) Fatty acid precursors of oxylipins. (B) Oxylipin subclasses monitored. (C) Extracted ion chromatogram of more polar oxylipins (prostaglandins, SPMs, leukotrienes, di-hydroxides, thromboxanes, lipoxins, and isoprostanes). (D) Extracted ion chromatogram of less polar oxylipins (mono-hydroxides, ketones, epoxides, and nitro fatty acids). Mass transitions used for extraction of ion chromatograms are described in Table 2. (E) Extracted ion chromatograms of HETE and EET isobaric species derived from AA. Proposed fragmentations for HETE isobaric species are also displayed. (F) Extracted ion chromatograms of HODE and EpOME isobaric species derived from LA. Specific fragments and proposed fragmentations for HODE isobaric species are also displayed. (G) MS/MS spectra, proposed fragmentations, and extracted ion chromatograms of 15-LXA4 and 6-LXA4. (H) MS/MS spectra, proposed fragmentations, and extracted ion chromatograms of PGE2 and PGD2. Abbreviations: OA, Oleic acid; LA, Linoleic acid; ALA, α-linolenic acid; GLA, γ-linolenic acid; EDA, eicosadienoic acid; MA, mead acid; DGLA, dihomo-γ-linolenic acid; AA, arachidonic acid; EPA, eicosapentaenoic acid; AdA, adrenic acid; DPA, docosapentaenoic acid; DHA, docosahexaenoic acid; IS, internal standard; SPMs, specialized pro-resolving mediators; HETE, hydroxy-eicosatetraenoic acid; EET, epoxy-eicosatrienoic acid; HODE, hydroxy-octadecadienoic acid; EpOME, epoxy-octadecenoic acid.

**Table 2.**
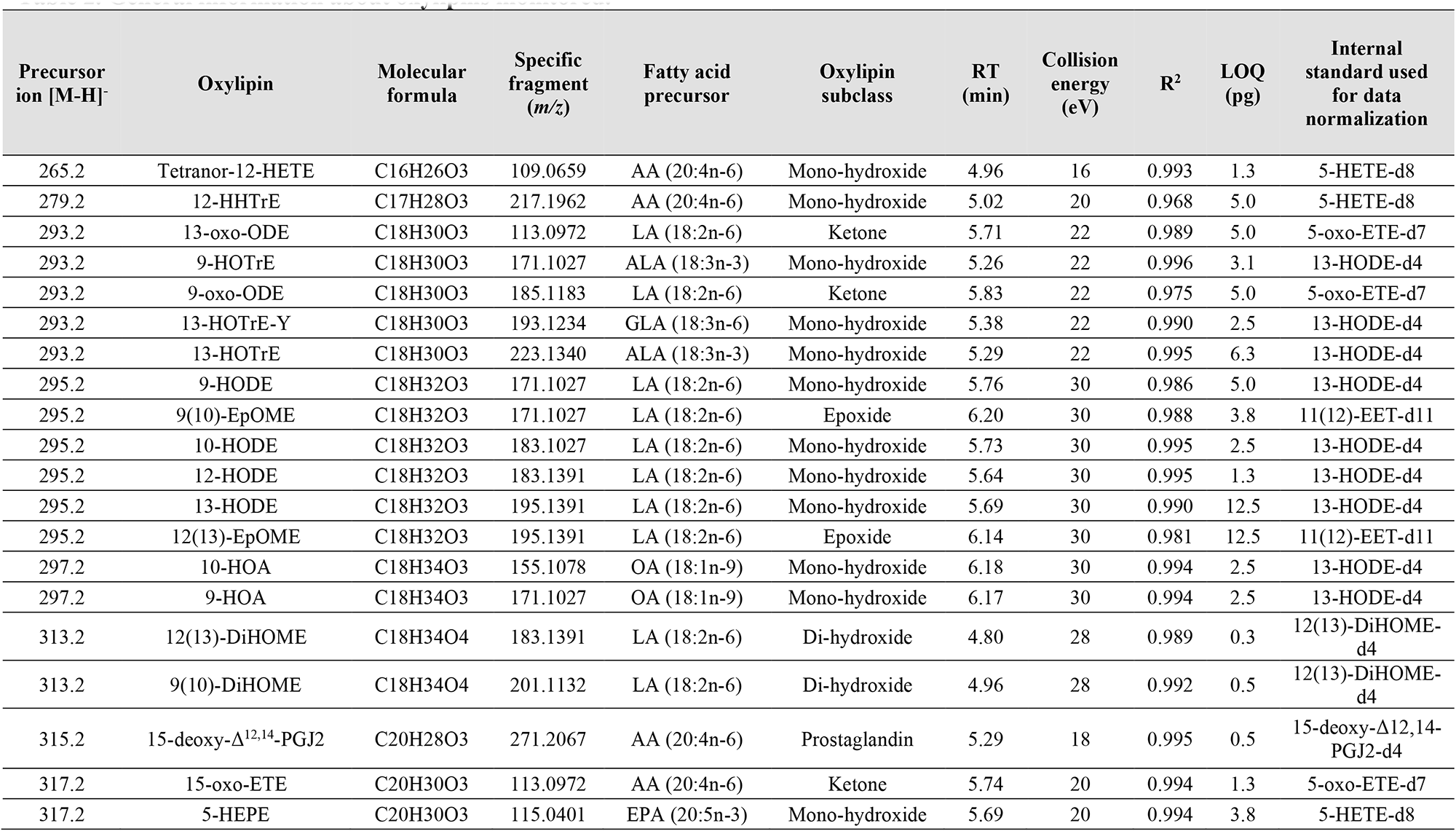
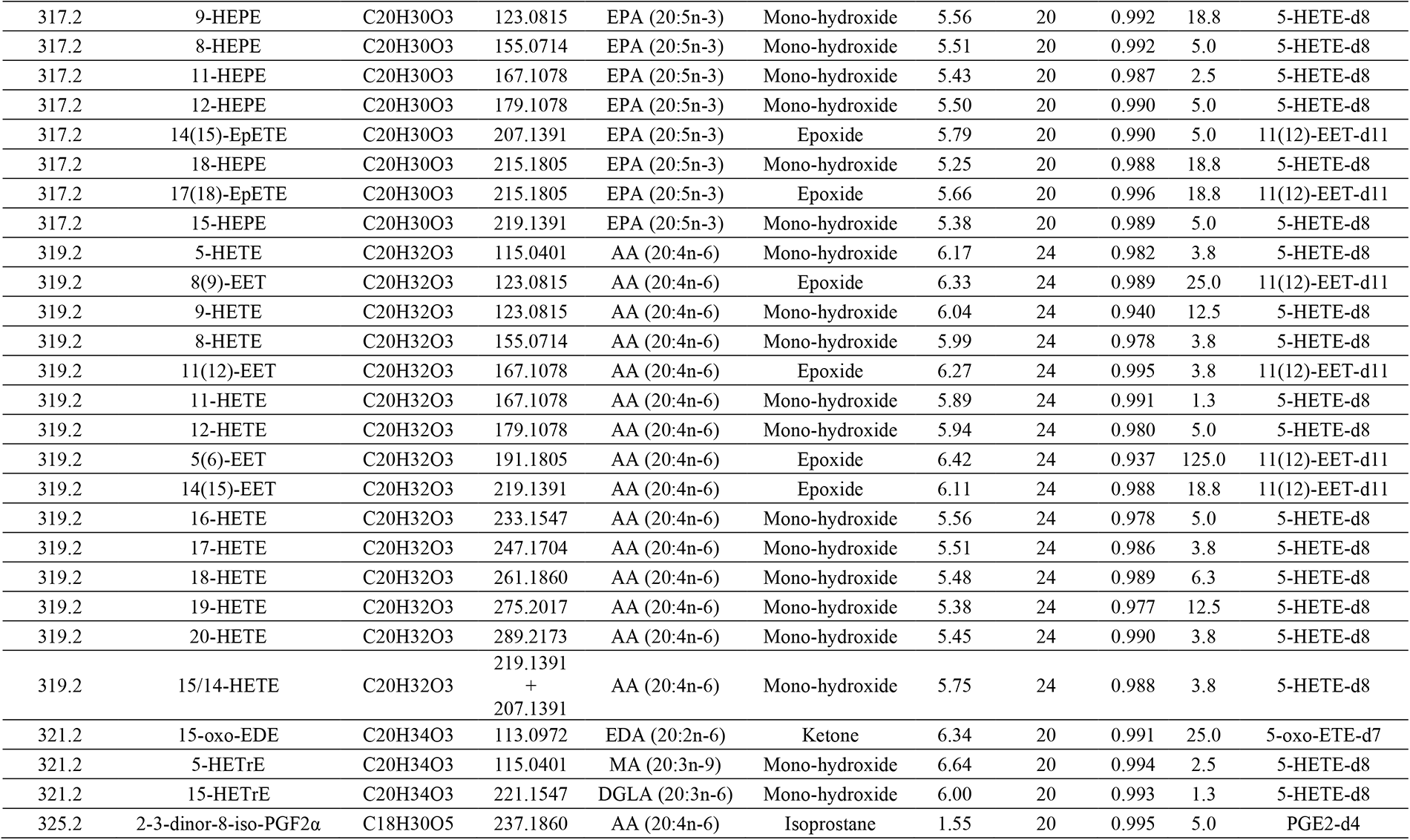
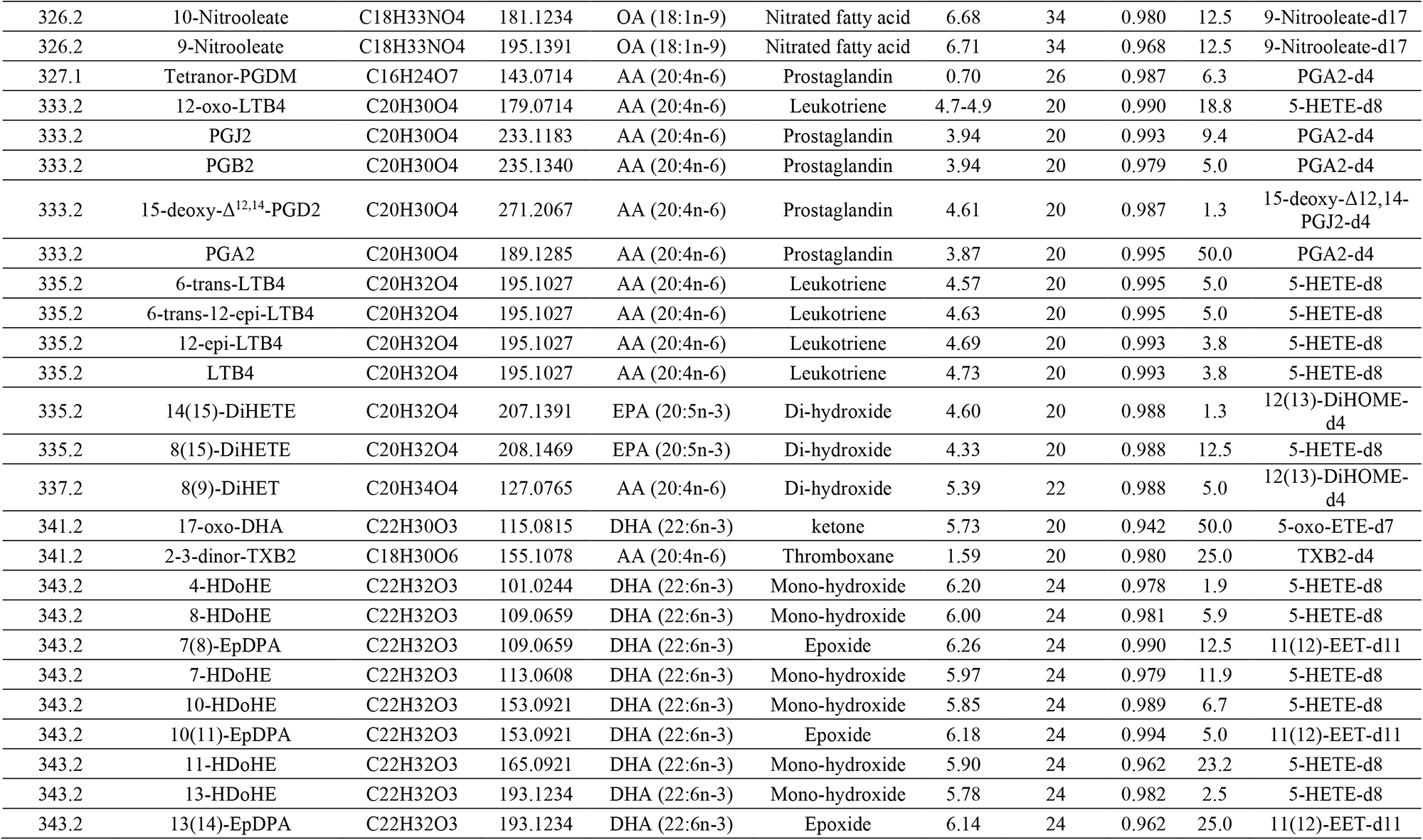
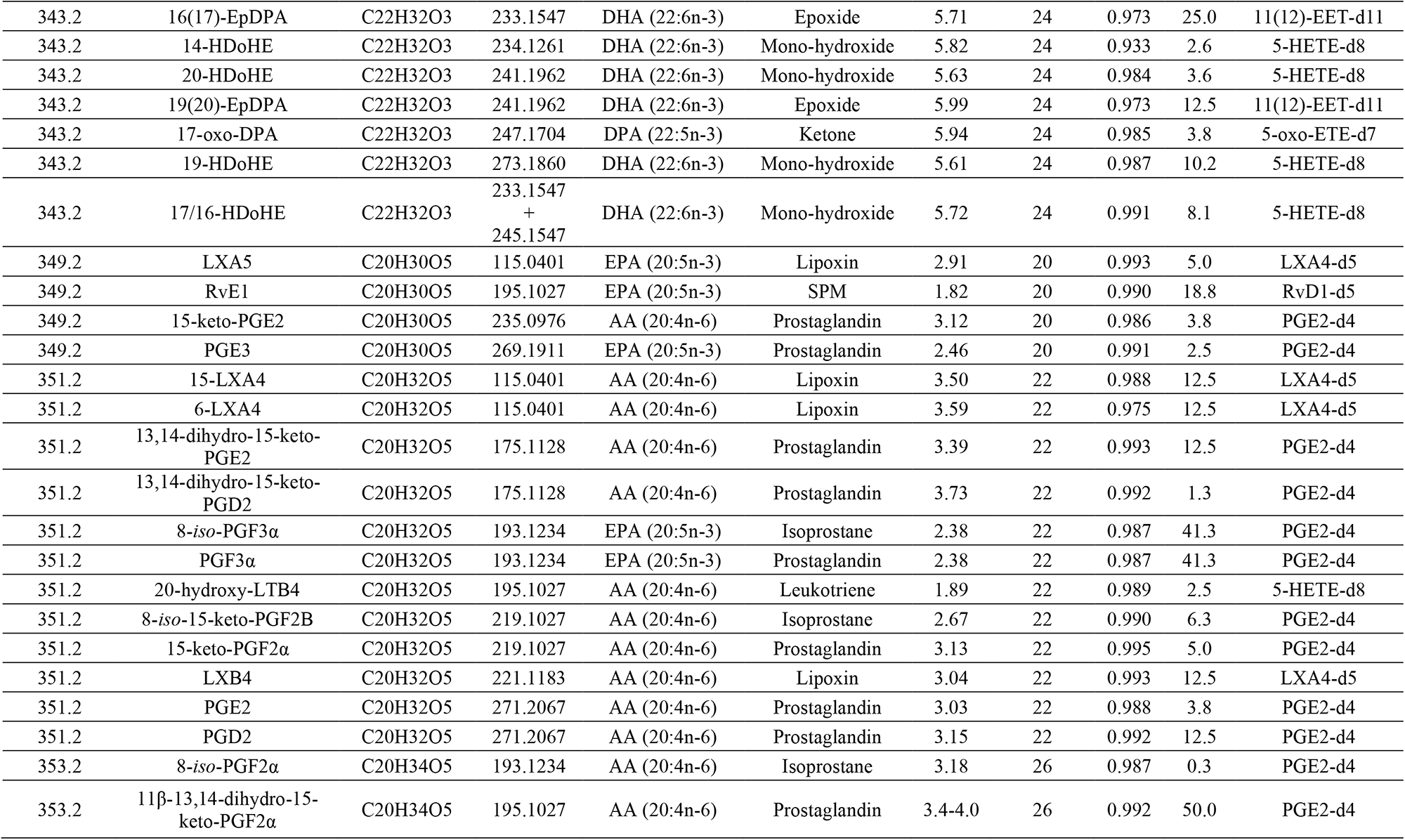
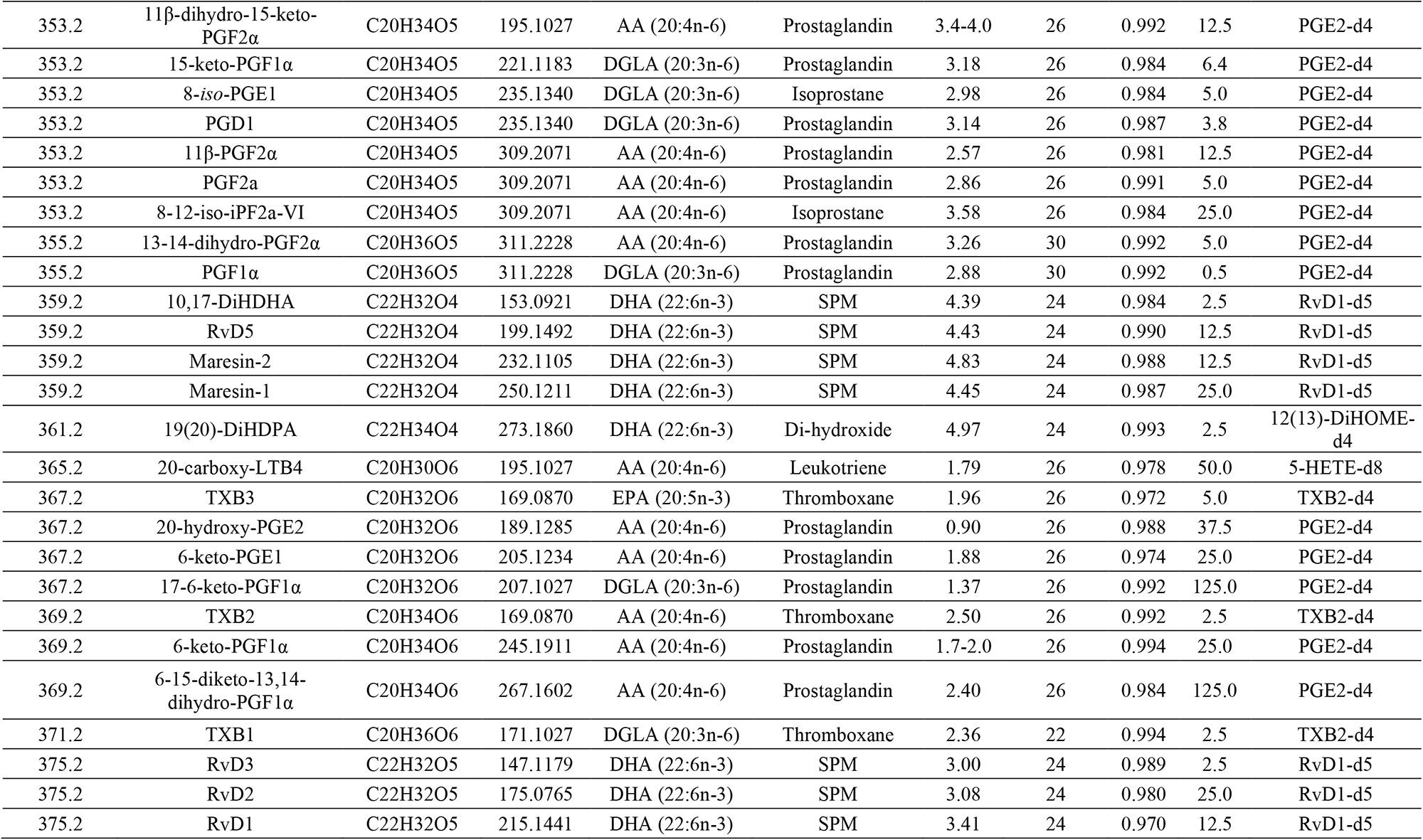
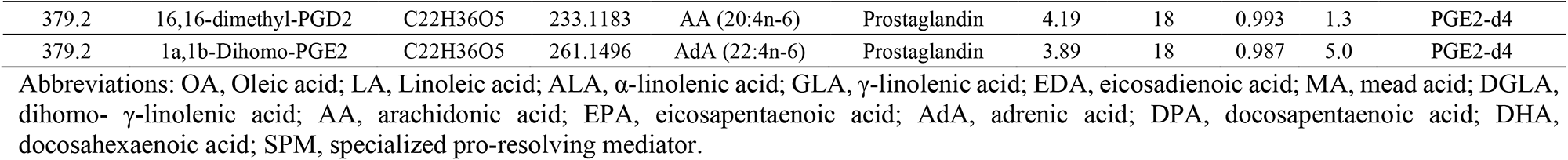
General information about oxylipins monitored.

Most oxylipins are present at very low concentration (pM to nM) in blood plasma and have high chemical similarity, and their precise identification and quantification remains an analytical challenge. Oxylipins frequently generate non-specific fragments, such as loss of carbon dioxide (CO_2_) and water (H_2_O) molecules, and therefore, contributing little for molecule identification. Moreover, the use of highly specific fragments generated at very low yields can negatively affect method sensitivity. Thus, to improve both the accuracy and sensitivity of their identification we performed a detailed inspection of the fragmentation profile of each oxylipin standard by direct infusion high resolution MS/MS analysis. Fragments were selected based on the criteria to be not only the most specific fragment as possible, but primarily the most intense fragment. In addition, we performed a collision energy ramp analysis of each analyte to assess the best fragmentation conditions for the specific fragments (Table 2; Fig. S2). Extracted ion chromatograms of polar oxylipins containing 2 or more oxygen atoms (excluding the ones in carboxyl group), such as prostaglandins, leukotrienes, lipoxins, and others show retention time (RT) between 1 and 6 min (Fig. 1C). On the other hand, non-polar oxylipins (1 oxygen atom), such as mono-hydroxides, ketones, and epoxides eluted after 5 min (Fig. 1D). Mass transitions used for extraction of ion chromatograms are described in Table 2. Once most of oxylipins are present as isobaric species, we highlighted the AA-derived mono-hydroxides, namely as hydroxy-eicosatetraenoic acids (HETEs), to emphasize the specificity of the fragments used for oxylipin analysis (Fig. 1E). Proposed fragmentations for HETE species are also displayed in Fig. 1E. Notably, the AA-derived epoxy-eicosatrienoic acids (EETs) show the same precursor ion ([M-H]^-^ = 319.2) and fragments as the HETEs. For instance, the 8(9)-EET (*m/z* 155.0714 or 123.0815) shows both specific fragments identified for the 8-HETE (*m/z* 155.0714) and 9-HETE (*m/z* 123.0815). However, EET and HETE isomers are easily differentiated according to their RT (Fig. 1E).

We also highlight the substantial contribution of LC-HRMS for accurate identification of oxylipins showing similar fragments. As depicted in Fig. 1F, the LA-derived mono-hydroxides, namely as hydroxy-octadecadienoic acid (HODEs), show 4 isobaric species. Notably, the specific fragments of 10-HODE (*m/z* 183.1027) and 12-HODE (*m/z* 183.1391) isomers can be only resolved by HRMS (Fig. 1F). Importantly, some isobaric oxylipin species show the same fragmentation profile. As displayed in Fig. 1G, both lipoxins 15(*R*)-LXA_4_ and 6(*S*)-LXA_4_ share the same fragmentation profile. However, chromatographic separation allows their differentiation according to the RT (15(*R*)-LXA_4_ = 3.50 min; 6(*S*)-LXA_4_ = 3.59 min) (Fig. 1G, Table 2). Similarly, both PGE_2_ and PGD_2_ share the same fragmentation profile (Fig. 1H). Thus, we were able to differentiate them exclusively according to their RT (PGE_2_ = 3.02 min; PGD_2_ = 3.15 min) (Fig. 1H, Table 2).

Following the method validation, we accessed the coefficient of determination (R^2^) and limit of quantification (LOQ) of each oxylipin. Data obtained from method validation show high linearity (R^2^ ≥ 0.980 for 82% of analytes) and sensitivity (LOQ between 0.3 and 25 pg for 92% of analytes) for most of the oxylipins (Table 2). The sensitivity achieved in our method shows the same order of magnitude to data obtained in low resolution hybrid (triple quadrupole / linear ion trap) mass spectrometer [26]. After adjustment of the chromatographic conditions for the LC-HRMS method, we constructed intra- and inter-day calibration curves. To proceed with method validation, we checked the precision and accuracy, which were accessed through the analysis of quality controls at high, medium, and low concentration of each oxylipin intra-(Table S2) and inter-daily (Table S3). Most quality controls showed coefficients of variation below 15% in the intra-(97% of measurements) and inter-day (93% of measurements) assays. In addition, the quality controls showed high accuracy (between 85 and 115%) as evidenced in the intra-(88% of measurements) and inter-day (86% of measurements) assays. To determine the oxylipin extraction efficiency using solid phase extraction (SPE), we checked the recovery rate of labelled internal standards spiked into high volume of blood plasma (500 µL) (Table 3). This analysis showed that all the 12 internal standards were extracted from plasma without major losses, being the recovery rate higher than 70% (Table 3). Regarding the matrix effect, 7 from 12 internal standards showed values above 70% (Table 3). Interestingly, we observed major interferences of matrix on internal standards showing retention time > 6 min, such as 5-HETE-d8, 5-oxo-ETE-d7, 11(12)-EET-d11, and 9-nitrooleate-d17 (Table 1). This finding may be linked with signal suppression caused by elution during the oxylipin sample extraction of highly abundant lipids in blood plasma, such as phosphatidylcholines (PCs) and lyso-phosphatidylcholines (LPCs) [33]. In summary, the developed LC-HRMS method showed not only high specificity for accurate identification of oxylipins, but also great sensitivity and reproducibility for simultaneous quantification of 126 analytes in a single 15-min run.

**Table 3.**
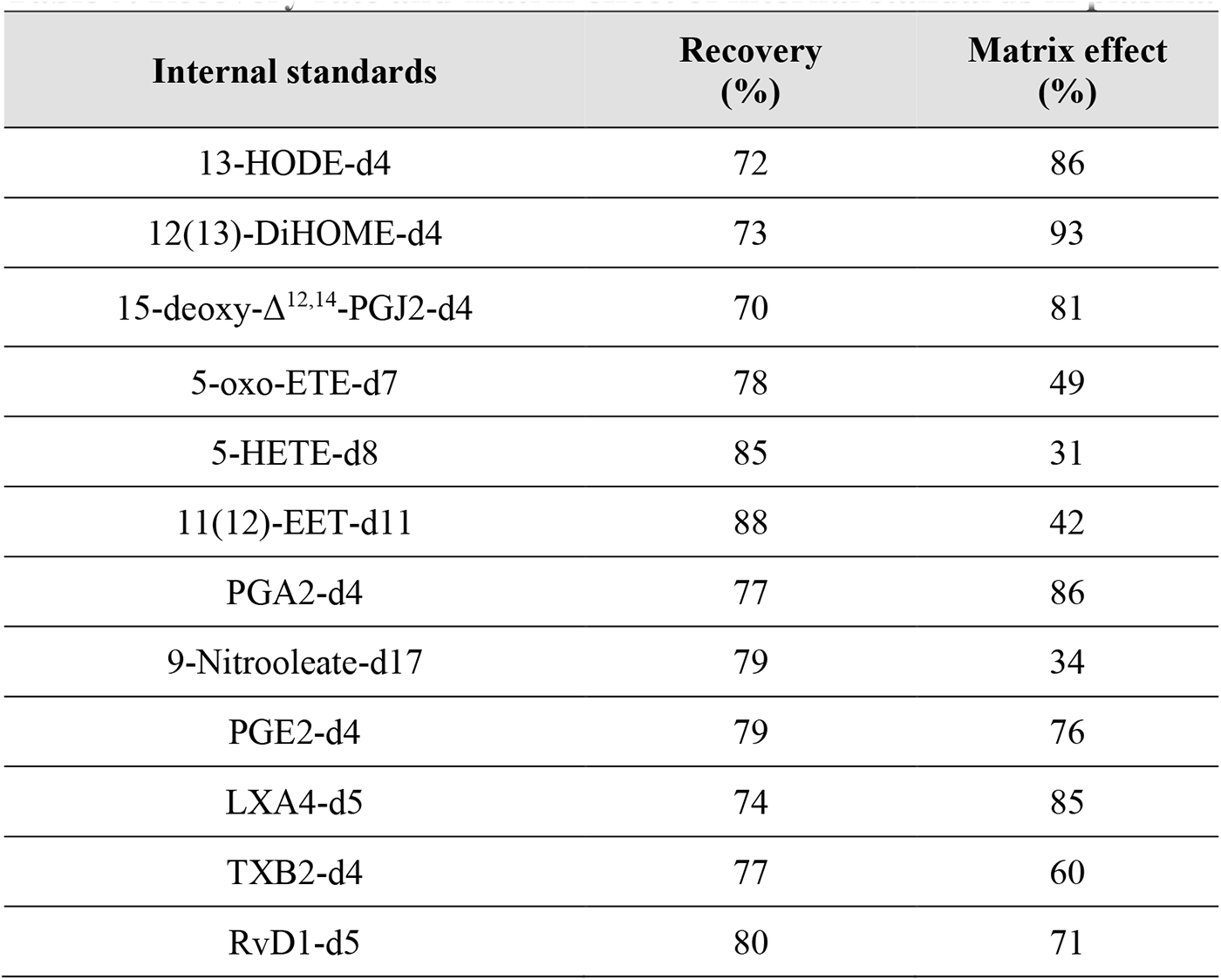
Recovery rate and matrix effect of internal standards in plasma.

### 3.2. Oxylipin analysis in plasma of ALS and WT rats revealed changes according to age and disease progression

To investigate how oxylipins are modulated in ALS, we applied the LC-HRMS method developed for analysis of oxylipins in blood plasma of asymptomatic (70 days) and symptomatic (120 days) ALS rats, and their respective age-matched wild type rats (WT 70 days and WT 120 days), as control groups. Here, we identified and quantified 56 oxylipins, which were sorted as lipid mono-hydroxides (35 species), di-hydroxides (4 species), ketones (3 species), prostaglandins (8 species), SPMs (3 species), and others (Fig. 2A; Table S4). The concentration average (expressed as log_2_ of ng/mL) of oxylipins in all experimental groups are displayed in the Fig. 2B-C. Notably, lipid mono-hydroxides derived from AA (5-HETE and 12-HETE), LA (9-HODE and 13-HODE) and DHA (4-HDoHE) were the most abundant oxylipins identified in plasma (Fig. 2B). In addition, less abundant oxylipins, such as prostaglandins (e.g., PGD2) and di-hydroxides (e.g., 6-trans-LTB4 and DiHOMEs), were potentially modulated in the experimental groups (Fig. 2C). Multivariate analysis performed by principal component analysis (PCA) showed a clear segregation between ALS 120d group and younger animals (WT 70d and ALS 70d groups), but a small segregation if compared to the WT 120d group (Fig. 2D). Interestingly, the spatial separation observed in PCA was mostly linked to alterations in lipid mono-hydroxides (12-HEPE, 12-HHTrE, 9-HETE, 9-HOA), di-hydroxides (9(10)- and 12(13)-DiHOMEs), thromboxanes (TXB2), and ketones (9-oxo-ODE, 13-oxo-ODE, and 15-oxo-ETE), as evidenced in the loadings plot of PCA (Fig. 2E). Univariate analysis performed by one-way ANOVA (p < 0.05; FDR adjusted) yielded 17 altered lipid molecular species when comparing all groups (Table S5). The altered oxylipins are displayed as clusters in the heatmap plot (Fig. 2F). Of note, most ALS 120d samples clustered together displaying increased levels of lipid mono-hydroxides, prostaglandins, lipoxins, ketones, among others. On the other hand, the content of two lipid di-hydroxides and one mono-hydroxide were decreased in ALS 120d group (Fig. 2F).

**Figure 2.**
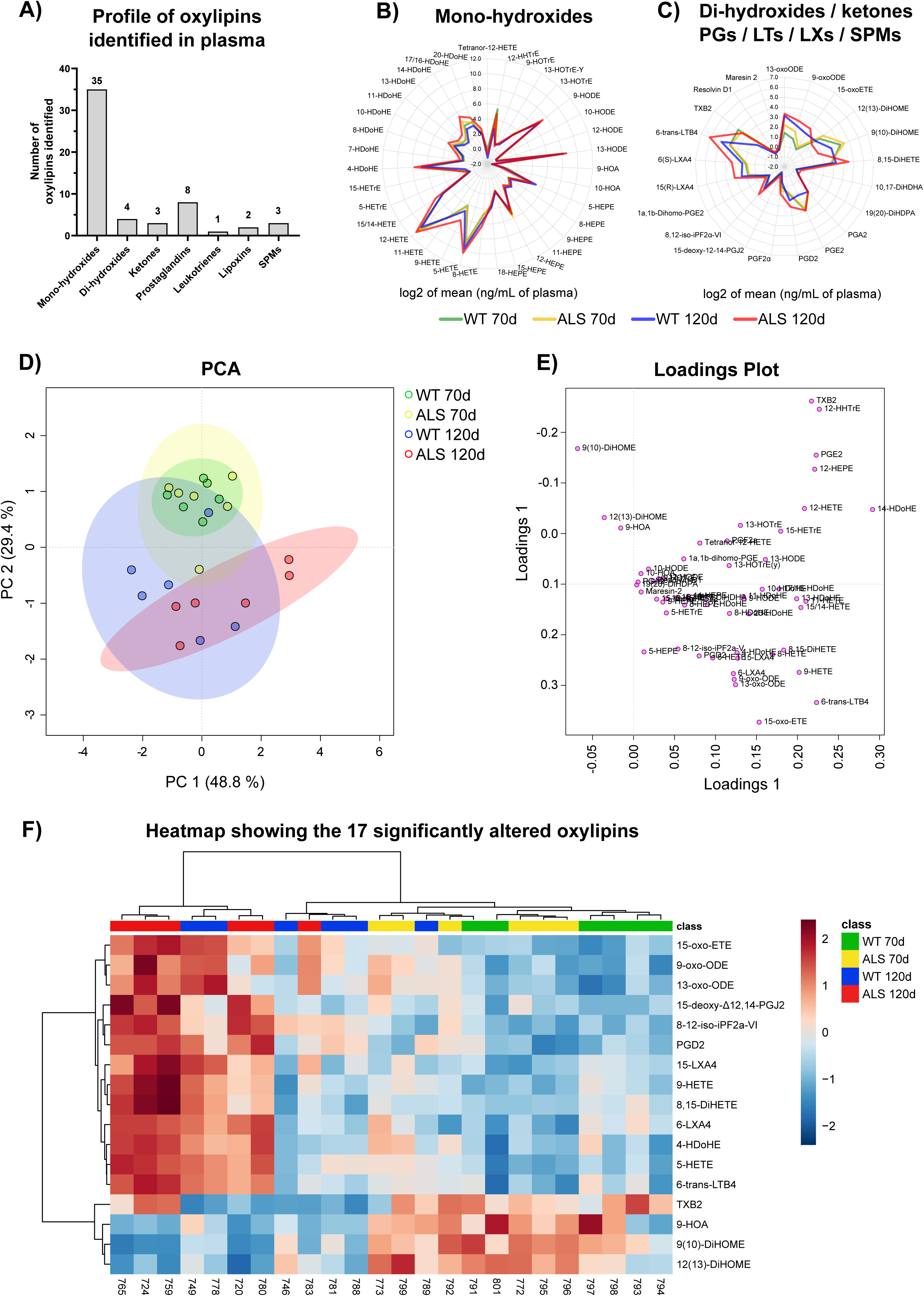
Univariate and multivariate analyses of oxylipins quantified in plasma suggest alterations according to ALS disease and age. (A) Profile of oxylipins identified in plasma of ALS and WT rats. (B) Concentration of lipid mono-hydroxides quantified in plasma of ALS and WT rats. (C) Concentration of lipid di-hydroxides, ketones, prostaglandins, leukotrienes, lipoxins and specialized pro-resolving mediators quantified in plasma of ALS and WT rats. Data (n=6) are shown as log_2_ of the average concentration (ng/mL of plasma) for each experimental group. (D) Principal component analysis (PCA) of oxylipin data. (E) Loadings plot of PCA. (F) Heatmap plot of oxylipins significantly altered in plasma of ALS rats. Data were log_10_ transformed prior to statistical analysis in Metaboanalyst. Heatmap plot displaying clusters of samples and the 17 significantly altered lipid molecular species according to one-way ANOVA followed by Tukey’s post-test (p < 0.05; FDR-adjusted). Individual lipids are shown in rows and samples displayed in columns, according to cluster analysis (clustering distance was calculated by Pearson and clustering algorithm estimated by Ward). Each colored cell on the heatmap plot corresponds to values above (red) or below (blue) the mean normalized concentrations for a given oxylipin.

Detailed inspection of alterations in ALS 120d group revealed increased content of several AA-derived oxylipins (Fig. 3). Indeed, such alterations were associated with increased oxidative stress in plasma of ALS 120d rats, as evidenced by the increase of a lipid mono-hydroxide (9-HETE) and isoprostane (8,12-*iso*-iPF2α-VI) (Fig. 3G-H). Interestingly, LOX-derived oxylipins (15-oxo-ETE, 6-trans-LTB4 and 5-HETE) were increased in both WT 120d and ALS 120d groups, as compared to the rats at 70 days old (Fig. 3A-B-E). Those alterations seem to be linked to age rather than the disease per se. Remarkably, we also observed increased content of pro-inflammatory prostaglandins (PGD_2_ and 15-deoxy-Δ^12,14^-PGJ_2_) in ALS 120d group, as compared to the other groups (Fig. 3J-K). PGD_2_ is one of the major COX products, and it readily undergoes dehydration to J_2_-series prostaglandins, such as 15-deoxy-^Δ12,14^-PGJ_2_ [34]. Interestingly, immunohistochemistry analysis revealed that 15-deoxy-^Δ12,14^-PGJ_2_ was increased in in motor neurons of sporadic ALS patients [35]. In addition, the same study reported that 15-deoxy-^Δ12,14^-PGJ_2_ induced apoptotic cell death of SH-SY5Y neuroblastoma cells [35]. Mechanistically, it was suggested that 15-deoxy-^Δ12,14^-PGJ_2_-induced apoptosis is dependent on p53 [35]. On the other hand, we observed increased content of lipoxins (6(*S*)-LXA_4_ and 15(*R*)-LXA_4_), which are described in the literature as pro-resolving mediators [4,36], in ALS 120d group (Fig. 3C-D). The increased content of lipoxins in plasma of ALS 120d rats may suggest that anti-inflammatory pathways are stimulated in response to acute inflammation present at a symptomatic stage. The biosynthetic pathway of both non-altered (written in blue) and significantly altered (written in red) AA-derived oxylipins are highlighted in Fig. 3L.

**Figure 3.**
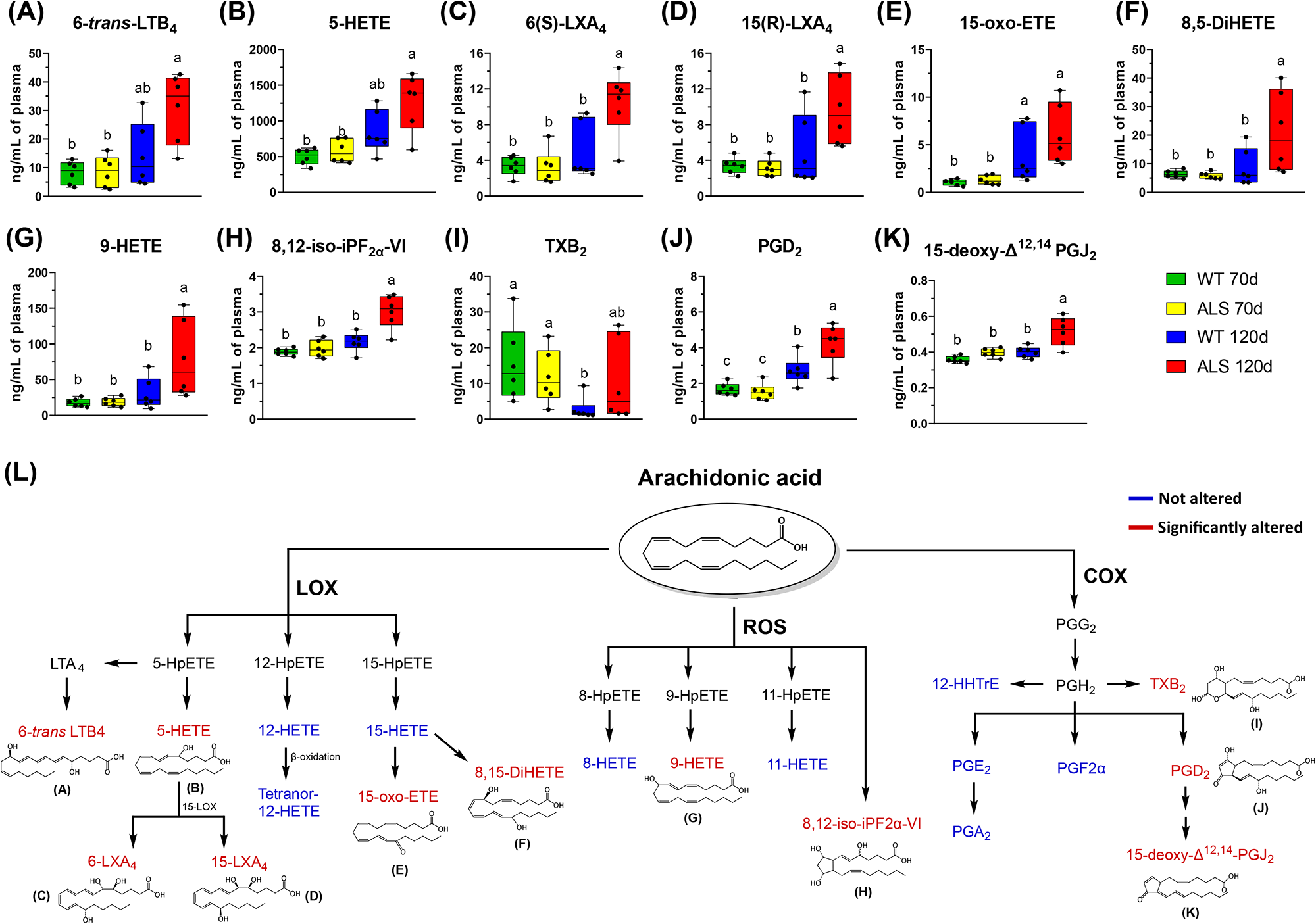
Oxylipins derived from AA are increased in plasma of ALS rats at symptomatic stage. (A-K) Significantly altered oxylipins derived from AA. Data (n = 6) are expressed as ng/mL of plasma and shown as box and whiskers. Means not sharing a common superscript are significantly different from each other. Statistical significance was evaluated by one-way ANOVA followed by Tukey’s post-test (p < 0.05; FDR-adjusted). (L) Scheme showing the not altered (written in blue) and significantly altered (written in red) AA-derived oxylipins. Chemical structures of altered oxylipins are also displayed. Abbreviations: LOX, lipoxygenase; ROS, reactive oxygen species; COX, cyclooxygenase; LTA_4_, Leukotriene A_4_; HpETE, hydroperoxy-eicosatetraenoic acid, PGG_2_, prostaglandin G_2_; PGH_2_, prostaglandin H_2_.

Although most of oxylipins altered in plasma of ALS 120d rats were derived from AA, we also observed important alterations in LA-derived oxylipins (Fig. 4). As depicted in Fig. 4A-B, both LA-derived ketones (9-oxo-ODE and 13-oxo-ODE) are increased in WT 120d ad ALS 120d groups, reinforcing the link of those alterations with age rather than the disease per se (Fig. 4A-B). Of note, 9-oxo-ODE and 13-oxo-ODE can be synthesized by hydroxy-fatty acids dehydrogenases from 9-HODE and 13-HODE, respectively. On the other hand, we observed drastic decrease of lipid di-hydroxides (9(10)-DiHOME and 12(13)-DiHOME) in rats at 120 days as compared to the groups at 70 days (Fig. 4C-D). Interestingly, the 9(10)-DiHOME was further decreased in ALS 120d group as compared to the other groups (Fig. 4C). DiHOMEs are generated from sequential oxidation of LA by cytochrome P450 (CYP) into epoxides (EpOMEs), followed by their hydrolysis into di-hydroxides, in a reaction catalyzed by the soluble epoxide hydrolase (sEH) (Fig. 4E) [37].

**Figure 4.**
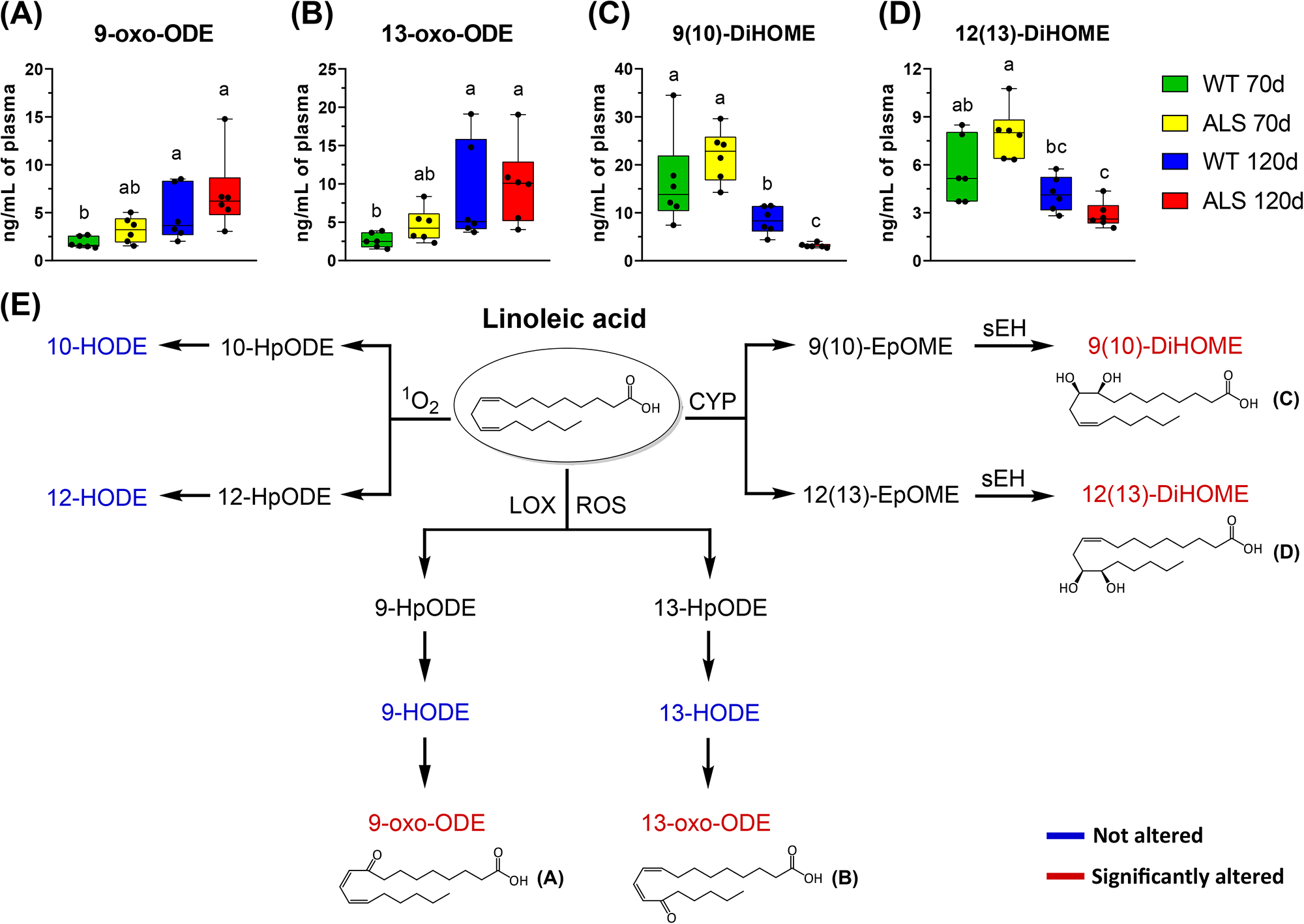
The Di-hydroxylated oxylipins derived from LA are decreased in plasma of ALS rats at symptomatic stage. (A-D) Significantly altered oxylipins derived from LA. Data (n = 6) are expressed as ng/mL of plasma and shown as box and whiskers. Means not sharing a common superscript are significantly different from each other. Statistical significance was evaluated by one-way ANOVA followed by Tukey’s post-test (p < 0.05; FDR-adjusted). (E) Scheme showing the not altered (written in blue) and significantly altered (written in red) LA-derived oxylipins. Chemical structures of altered oxylipins are also displayed. Abbreviations: ^1^O_2_, singlet oxygen; LOX, lipoxygenase; ROS, reactive oxygen species; CYP, cytochrome P450; sEH, soluble epoxide hydrolase; HpODE, hydroperoxy-octadecadienoic acid; HODE, hydroxy-octadecadienoic acid; oxo-ODE, oxo-octadecadienoic acid; EpOME, epoxy-octadecenoic acid; DiHOME, dihydroxy-octadecenoic acid.

To better understand the interaction among the oxylipins identified in plasma of ALS rats, we also performed a correlation analysis (Fig. 5). Correlation heatmap shows that lipid mono-hydroxides derived from AA (HETEs) and DHA (HDoHEs) are positively correlated (light red cluster in the center of heatmap) (Fig. 5A). On the other hand, DiHOMEs are negatively correlated with prostaglandins, HETEs, HDoHEs lipoxins, and ketones (light green cluster in the top of heatmap) (Fig. 5A). Correlation data and *p*-values are described in the Tables S6 and S7, respectively. Detailed inspection of the top 25 oxylipins correlated with 9(10)-and 12(13)-DiHOMEs confirmed that prostaglandins (PGD2 and 15-deoxy-PGJ2), isoprostane (8,12-iso-iPF2a-VI), lipoxins (15(*R*)-LXA4, 6(*S*)-LXA4) and AA-derived mono-hydroxides (5-HETE and 9-HETE) are strong and negatively connected with them. Interestingly, two OA-derived mono-hydroxides (9-HOA and 10-HOA) were positively correlated with 9(10)-DiHOME. Although there is no clear evidence in the literature about the mechanisms involved in their generation, it is likely that both 9-HOA and 10-HOA are oxidized by oxygenases, once OA is much less susceptible to undergo auto-oxidation, as compared to PUFAs [6].

**Figure 5.**
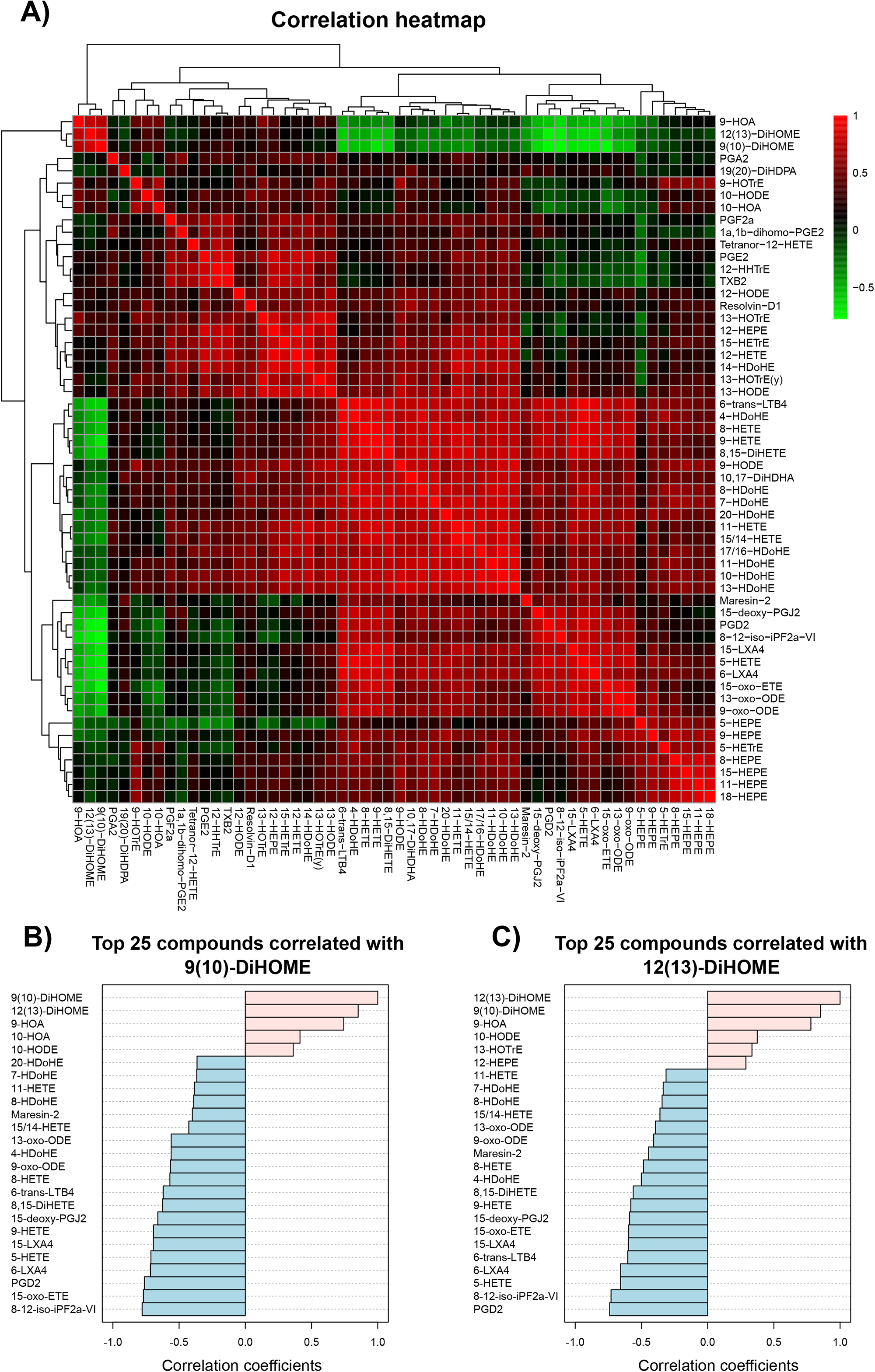
Correlation analysis shows that DiHOMEs are inversely correlated with AA-derived oxylipins in plasma of ALS rats. (A) Correlation analysis performed with oxylipin data. Clustering distance was calculated by Pearson r. Positive correlations are displayed in red, whereas the negative correlations are displayed in green. (B) Pattern analysis showing the top 25 compounds correlated with 9(10)-DiHOME. (C) Pattern analysis showing the top 25 compounds correlated with 9(10)-DiHOME. Correlation analysis was calculated by Pearson r.

Data from literature showed that 12(13)-DiHOME increases in response to cold exposure, and it is mostly synthesized in brown adipose tissue (BAT) [38]. This lipokine is described as a regulator of uptake of free fatty acids in BAT [38]. Mechanistically, it is suggested that 12(13)-DiHOME promotes the transfer of fatty acid transporters (CD36 and FATP1) from cytosol to plasma membrane [38]. In another study, it was demonstrated that the 12(13)-DiHOME released from BAT to the blood plasma during intensive exercise upregulates fatty acid uptake and β-oxidation in skeletal muscle [39]. ALS is frequently associated to lipid hypermetabolism, which leads to severe decrease in weight loss at a final stage of disease [40]. Indeed, it was suggested that fat content is a predictor of survival of ALS patients [41,42]. Thus, our data may support evidence that increased content of plasmatic DiHOMEs in ALS 70d rats is linked to lipid hypermetabolism at an early disease stage. This event is followed by a severe decrease of DiHOMEs ALS 120d rats, supporting the severe lipoatrophy observed in ALS at a symptomatic stage. In line with our findings, it was observed that oxylipins derived from LA, especially the HODEs and DiHOMEs were strongly downregulated in plasma of ALS patients [9].

In summary, we developed a robust method for global analysis of oxylipins by high resolution MS and applied it in an ALS rat model. The data obtained from this analysis revealed an increase of AA-derived oxylipins in plasma of ALS rats, which are mostly linked to inflammatory processes and oxidative stress, both strongly associated to ALS. On the other hand, we observed a decreased content of lipokines, namely DiHOMEs, which may be linked to hypermetabolism in ALS [43]. Taken together, our data support evidence that oxylipins can provide valuable information about the redox and metabolic status in ALS and support the investigation of new molecular mechanisms associated with the alterations observed in oxylipin metabolism.

## Acknowledgments

This work was supported by Fundação de Amparo à Pesquisa do Estado de São Paulo [FAPESP, CEPID-Redoxoma 2013/07937-8 and 10/50891-0]; Conselho Nacional de Desenvolvimento Científico e Tecnológico [CNPq, Universal 424094/2016-9 and PQ 309083/2017-6 and 313926/2021-2]; NAP-Redoxoma; Pró-Reitoria de Pesquisa USP; and CAPES. IG is supported by grant from FAPESP 2018/18633-3. A.B.C-F, R.S.S., I.F.D.P, A.I., and R.L.F are recipient of FAPESP scholarships.

## Author contributions

A.B.C-F. and L.S.Diniz performed the oxylipin method validation. A.B.C-F., H.O., I.F.D.P., L.S.Dantas., A.I., and R.L.F conducted the experiments with animals. A.B.C-F., R.S.S., and R.S.L. performed the oxylipin analysis in plasma. M.H.G.M. provided the SOD1-G93A rats for experiments. A.B.C-F. and I.G. performed the statistical analysis. A.B.C-F. and S.M. wrote the original draft. MYY and SM supervised the study. WTF and SM were responsible for funding acquisition. All authors edited and reviewed the final version of the manuscript.

## References

[1] T. Harayama, H. Riezman, Understanding the diversity of membrane lipid composition, Nat. Rev. Mol. Cell Biol. 19 (2018) 281–296. https://doi.org/10.1038/nrm.2017.138.

[2] G. Schmitz, J. Ecker, The opposing effects of n-3 and n-6 fatty acids, Prog. Lipid Res. 47 (2008) 147–155. https://doi.org/10.1016/j.plipres.2007.12.004.

[3] D. Wang, R.N. Dubois, Eicosanoids and cancer, Nat. Rev. Cancer. 10 (2010) 181–193. https://doi.org/10.1038/nrc2809.

[4] C.N. Serhan, N. Chiang, T.E. Van Dyke, Resolving inflammation: Dual anti-inflammatory and pro-resolution lipid mediators, Nat. Rev. Immunol. 8 (2008) 349–361. https://doi.org/10.1038/nri2294.

[5] T. Finkel, N.J. Holbrook, Oxidants, oxidative stress and the biology of ageing, Nature. 408 (2000) 239–247. https://doi.org/10.1038/35041687.

[6] H. Yin, L. Xu, N.A. Porter, Free Radical Lipid Proxidation: Mechanisms and Analysis., Chem. Rev. 111 (2011) 5944–5972. https://doi.org/10.1021/cr200084z.

[7] N. Chiang, C.N. Serhan, Molecular Aspects of Medicine Structural elucidation and physiologic functions of specialized pro-resolving mediators and their receptors, Mol. Aspects Med. 58 (2017) 114–129. https://doi.org/10.1016/j.mam.2017.03.005.

[8] D.S. Straus, C.K. Glass, Cyclopentenone prostaglandins: New insights on biological activities and cellular targets, Med. Res. Rev. 21 (2001) 185–210. https://doi.org/10.1002/med.1006.

[9] M. Mastrogiovanni, A. Trostchansky, H. Naya, R. Dominguez, C. Marco, M. Povedano, R. López-Vales, H. Rubbo, HPLC-MS/MS Oxylipin Analysis of Plasma from Amyotrophic Lateral Sclerosis Patients, Biomedicines. 10 (2022). https://doi.org/10.3390/BIOMEDICINES10030674.

[10] N.G. Bazan, V. Colangelo, W.J. Lukiw, Prostaglandins and other lipid mediators in Alzheimer’s disease, Prostaglandins Other Lipid Mediat. 68–69 (2002). https://doi.org/10.1016/S0090-6980(02)00031-X.

[11] O. Hardiman, Z.S., Ammar Al-Chalabi, Adriano, Chio, Emma M. Corr, Giancarlo Logroscino, Wim Robberecht, Pamela J. Shaw, and L.H. van den Berg, Amyotrophic lateral sclerosis, Nat. Rev. | Dis. Prim. 3 (2017) 1–18. https://doi.org/10.1201/b13434.

[12] J.K. Andersen, Oxidative stress in neurodegeneration: Cause or consequence?, Nat. Rev. Neurosci. 10 (2004) S18. https://doi.org/10.1038/nrn1434.

[13] B. Halliwell, Oxidative stress and neurodegeneration: Where are we now?, J. Neurochem. 97 (2006) 1634–1658. https://doi.org/10.1111/j.1471-4159.2006.03907.x.

[14] A.B. Chaves-Filho, I.F.D. Pinto, L.S. Dantas, A.M. Xavier, A. Inague, R.L. Faria, M.H.G. Medeiros, I. Glezer, M.Y. Yoshinaga, S. Miyamoto, Alterations in lipid metabolism of spinal cord linked to amyotrophic lateral sclerosis, Sci. Rep. 9 (2019) 1–14. https://doi.org/10.1038/s41598-019-48059-7.

[15] A. Trostchansky, M. Mastrogiovanni, E. Miquel, S. Rodríguez-Bottero, L. Martínez-Palma, P. Cassina, H. Rubbo, Profile of arachidonic acid-derived inflammatory markers and its modulation by nitro-oleic acid in an inherited model of amyotrophic lateral sclerosis, Front. Mol. Neurosci. 11 (2018). https://doi.org/10.3389/fnmol.2018.00131.

[16] H. Miyagishi, Y. Kosuge, A. Takano, M. Endo, H. Nango, S. Yamagata-Murayama, D. Hirose, R. Kano, Y. Tanaka, K. Ishige, Y. Ito, Increased Expression of 15-Hydroxyprostaglandin Dehydrogenase in Spinal Astrocytes During Disease Progression in a Model of Amyotrophic Lateral Sclerosis, Cell. Mol. Neurobiol. 37 (2017). https://doi.org/10.1007/s10571-016-0377-9.

[17] G. Almer, P. Teismann, Z. Stevic, J. Halaschek-Wierner, L. Deecke, V. Kostic, S. Przedborski, Increased levels of the pro-inflammatory prostaglandin PGE2 in CSF from ALS patients, Neurology. 58 (2002) 1277–1279.

[18] P. Bezzi, G. Carmignoto, L. Pasti, S. Vesce, D. Rossi, B.L. Rizzini, T. Pozzant, A. Volterra, Prostaglandins stimulate calcium-dependent glutamate release in astrocytes, Nature. 391 (1998) 281–285. https://doi.org/10.1038/34651.

[19] X. Liang, Q. Wang, J. Shi, L. Lokteva, R.M. Breyer, T.J. Montine, K. Andreasson, The prostaglandin E2 EP2 receptor accelerates disease progression and inflammation in a model of amyotrophic lateral sclerosis, Ann. Neurol. 64 (2008) 304–314. https://doi.org/10.1002/ANA.21437.

[20] M. Bilak, L. Wu, Q. Wang, N. Haughey, K. Conant, C. St. Hillaire, K. Andreasson, PGE2 receptors rescue motor neurons in a model of amyotrophic lateral sclerosis, Ann. Neurol. 56 (2004) 240–248. https://doi.org/10.1002/ANA.20179.

[21] O. Quehenberger, A.M. Armando, A.H. Brown, S.B. Milne, D.S. Myers, A.H. Merrill, S. Bandyopadhyay, K.N. Jones, S. Kelly, R.L. Shaner, C.M. Sullards, E. Wang, R.C. Murphy, R.M. Barkley, T.J. Leiker, C.R.H. Raetz, Z. Guan, G.M. Laird, D.A. Six, D.W. Russell, J.G. McDonald, S. Subramaniam, E. Fahy, E.A. Dennis, Lipidomics reveals a remarkable diversity of lipids in human plasma, J. Lipid Res. 51 (2010) 3299–3305. https://doi.org/10.1194/JLR.M009449.

[22] M.R. Wenk, Lipidomics: New tools and applications, Cell. 143 (2010) 888–895. https://doi.org/10.1016/j.cell.2010.11.033.

[23] A. Shevchenko, K. Simons, Lipidomics: Coming to grips with lipid diversity, Nat. Rev. Mol. Cell Biol. 11 (2010) 593–598. https://doi.org/10.1038/nrm2934.

[24] B. Burla, M. Arita, M. Arita, A.K. Bendt, A. Cazenave-Gassiot, E.A. Dennis, K. Ekroos, X. Han, K. Ikeda, G. Liebisch, M.K. Lin, T.P. Loh, P.J. Meikle, M. Orešič, O. Quehenberger, A. Shevchenko, F. Torta, M.J.O. Wakelam, C.E. Wheelock, M.R. Wenk, MS-based lipidomics of human blood plasma: A community-initiated position paper to develop accepted guidelines, J. Lipid Res. 59 (2018) 2001–2017. https://doi.org/10.1194/jlr.S087163.

[25] A. Kij, K. Kus, I. Czyzynska-Cichon, S. Chlopicki, M. Walczak, Development and validation of a rapid, specific and sensitive LC-MS/MS bioanalytical method for eicosanoid quantification – assessment of arachidonic acid metabolic pathway activity in hypertensive rats, Biochimie. 171–172 (2020). https://doi.org/10.1016/j.biochi.2020.03.010.

[26] Y. Wang, A.M. Armando, O. Quehenberger, C. Yan, E.A. Dennis, Comprehensive ultra-performance liquid chromatographic separation and mass spectrometric analysis of eicosanoid metabolites in human samples, J Chromatogr A. 1359 (2014) 60–69. https://doi.org/10.1016/j.chroma.2014.07.006.

[27] P.B.M.C. Derogis, F.P. Freitas, A.S.F. Marques, D. Cunha, P.P. Appolinário, F. de Paula, T.C. Lourenço, M. Murgu, P. Di Mascio, M.H.G. Medeiros, S. Miyamoto, The Development of a Specific and Sensitive LC-MS-Based Method for the Detection and Quantification of Hydroperoxy- and Hydroxydocosahexaenoic Acids as a Tool for Lipidomic Analysis, PLoS One. 8 (2013) 1–14. https://doi.org/10.1371/journal.pone.0077561.

[28] S. Gouveia-Figueira, M.L. Nording, Validation of a tandem mass spectrometry method using combined extraction of 37 oxylipins and 14 endocannabinoid-related compounds including prostamides from biological matrices, Prostaglandins Other Lipid Mediat. 121 (2015). https://doi.org/10.1016/j.prostaglandins.2015.06.003.

[29] S. Miyamoto, G.R. Martinez, A.P.B. Martins, M.H.G. Medeiros, P. Di Mascio, Direct evidence of singlet molecular oxygen [O2 (1Δg)] production in the reaction of linoleic acid hydroperoxide with peroxynitrite, J. Am. Chem. Soc. 125 (2003) 4510–4517. https://doi.org/10.1021/ja029262m.

[30] C.A. Sorgi, A.P.F. Peti, T. Petta, A.F.G. Meirelles, C. Fontanari, L.A.B. de Moraes, L.H. Faccioli, Data descriptor: Comprehensive high-resolution multiple-reaction monitoring mass spectrometry for targeted eicosanoid assays, Sci. Data. 5 (2018). https://doi.org/10.1038/sdata.2018.167.

[31] C. Hinz, S. Liggi, G. Mocciaro, S. Jung, I. Induruwa, M. Pereira, C.E. Bryant, S.W. Meckelmann, V.B. O’Donnell, R.W. Farndale, J. Fjeldsted, J.L. Griffin, A Comprehensive UHPLC Ion Mobility Quadrupole Time-of-Flight Method for Profiling and Quantification of Eicosanoids, Other Oxylipins, and Fatty Acids, Anal. Chem. 91 (2019). https://doi.org/10.1021/acs.analchem.8b04615.

[32] S. Hellhake, S.W. Meckelmann, M.T. Empl, K. Rentmeister, W. Wißdorf, P. Steinberg, O.J. Schmitz, T. Benter, N.H. Schebb, Non-targeted and targeted analysis of oxylipins in combination with charge-switch derivatization by ion mobility high-resolution mass spectrometry, Anal. Bioanal. Chem. 412 (2020). https://doi.org/10.1007/s00216-020-02795-2.

[33] O.A. Ismaiel, M.S. Halquist, M.Y. Elmamly, A. Shalaby, H.T. Karnes, Monitoring phospholipids for assessment of matrix effects in a liquid chromatography-tandem mass spectrometry method for hydrocodone and pseudoephedrine in human plasma, J. Chromatogr. B. Analyt. Technol. Biomed. Life Sci. 859 (2007) 84–93. https://doi.org/10.1016/J.JCHROMB.2007.09.007.

[34] Y. Kikawa, S. Narumiya, M. Fukushima, H. Wakatsuka, O. Hayaishi, 9-Deoxy-delta 9, delta 12-13,14-dihydroprostaglandin D2, a metabolite of prostaglandin D2 formed in human plasma, Proc. Natl. Acad. Sci. U. S. A. 81 (1984) 1317– 1321. https://doi.org/10.1073/PNAS.81.5.1317.

[35] M. Kondo, T. Shibata, T. Kumagai, T. Osawa, N. Shibata, M. Kobayashi, S. Sasaki, M. Iwata, N. Noguchi, K. Uchida, 15-Deoxy-Δ12,14-prostaglandin J2: The endogenous electrophile that induces neuronal apoptosis, Proc. Natl. Acad. Sci. U. S. A. 99 (2002). https://doi.org/10.1073/pnas.112212599.

[36] D. Press, Lipoxins: nature’s way to resolve inflammation, J. Inflamm. Res. (2015) 181–192.

[37] K. Hildreth, S.D. Kodani, B.D. Hammock, L. Zhao, Cytochrome P450-derived linoleic acid metabolites EpOMEs and DiHOMEs: a review of recent studies, J. Nutr. Biochem. 86 (2020). https://doi.org/10.1016/J.JNUTBIO.2020.108484.

[38] M.D. Lynes, L.O. Leiria, M. Lundh, A. Bartelt, F. Shamsi, T.L. Huang, H. Takahashi, M.F. Hirshman, C. Schlein, A. Lee, L.A. Baer, F.J. May, F. Gao, N.R. Narain, E.Y. Chen, M.A. Kiebish, A.M. Cypess, M. Blüher, L.J. Goodyear, G.S. Hotamisligil, K.I. Stanford, Y. Tseng, The cold-induced lipokine 12,13-diHOME promotes fatty acid transport into brown adipose tissue, Nat. Med. 23 (2017). https://doi.org/10.1038/nm.4297.

[39] K.I. Stanford, M.D. Lynes, H. Takahashi, Y. Tseng, P.M. Coen, L.J. Goodyear, K.I. Stanford, M.D. Lynes, H. Takahashi, L.A. Baer, P.J. Arts, F.J. May, 12,13-diHOME: An Exercise-Induced Lipokine that Increases Skeletal Muscle Fatty Acid Uptake, Cell Metab. 27 (2018) 1111–1120.e3. https://doi.org/10.1016/j.cmet.2018.03.020.

[40] C. Moglia, A. Calvo, M. Grassano, A. Canosa, U. Manera, F. D’ovidio, A. Bombaci, E. Bersano, L. Mazzini, G. Mora, A. Chiò, S. Cammarosano, R. Vasta, M.C. Torrieri, L. Solero, M. Clerico, S. De Mercanti, A. Mauro, L. Pradotto, F. De Marchi, L. Sosso, D. Leotta, L. Appendino, D. Imperiale, R. Cavallo, C. Geda, F. Poglio, P. Santimaria, U. Massazza, A. Villani, R. Conti, L.C. Ruiz, M. Palermo, F. Vergnano, E. Rota, M.T. Penza, M. Aguggia, P. Meineri, P. Ghiglione, N. Launaro, G. Astegiano, G. Corso, Early weight loss in amyotrophic lateral sclerosis: outcome relevance and clinical correlates in a population-based cohort, J. Neurol. Neurosurg. Psychiatry. 90 (2019) 666–673. https://doi.org/10.1136/JNNP-2018-319611.

[41] I. Lee, M. Kazamel, T. McPherson, J. McAdam, M. Bamman, A. Amara, D.L. Smith, P.H. King, Fat mass loss correlates with faster disease progression in amyotrophic lateral sclerosis patients: Exploring the utility of dual-energy x-ray absorptiometry in a prospective study, PLoS One. 16 (2021) e0251087. https://doi.org/10.1371/JOURNAL.PONE.0251087.

[42] E. Lindauer, L. Dupuis, H.P. Müller, H. Neumann, A.C. Ludolph, J. Kassubek, Adipose Tissue Distribution Predicts Survival in Amyotrophic Lateral Sclerosis, PLoS One. 8 (2013). https://doi.org/10.1371/JOURNAL.PONE.0067783.

[43] L. Dupuis, J.-P. Loeffl, L. Dupuis, P.-F. Pradat, A.C. Ludolph, J.-P. Loeffl, Energy metabolism in amyotrophic lateral sclerosis, Lancet Neurol. 10 (2011) 75–82. https://doi.org/10.1016/S1474.

