## Supplemental Figures for "Plasma Oxylipin Profiling by High Resolution Mass Spectrometry Reveal Signatures of Inflammation and Hypermetabolism in Amyotrophic Lateral Sclerosis"

**Supplementary figures**

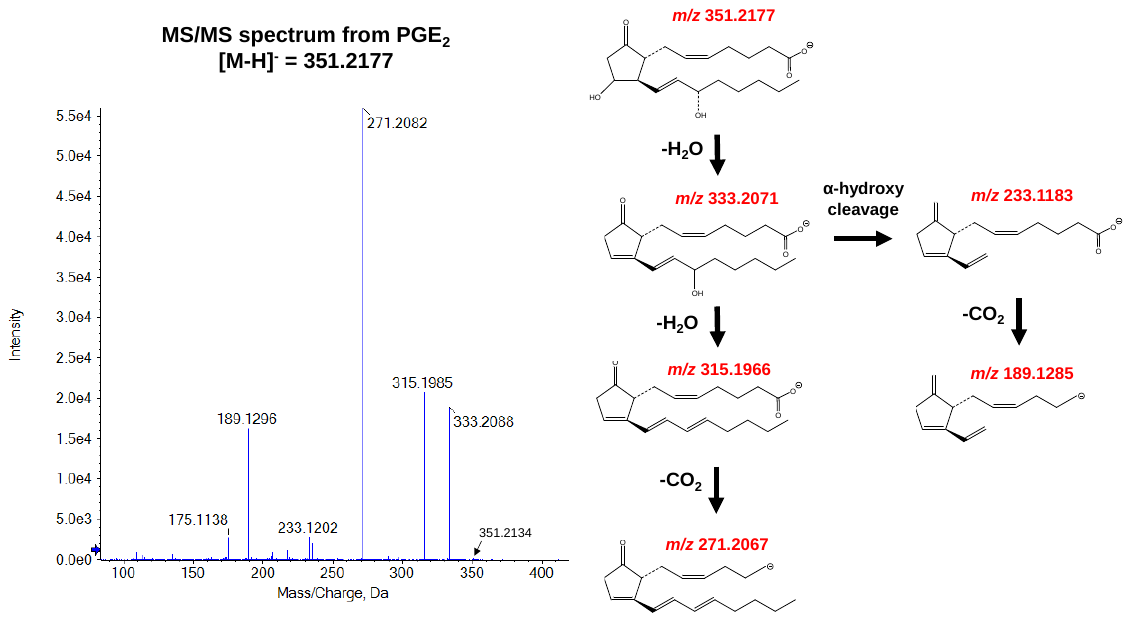

**Figure S1. Representative MS/MS spectrum of PGE_2_ and chemical structures of fragments.**

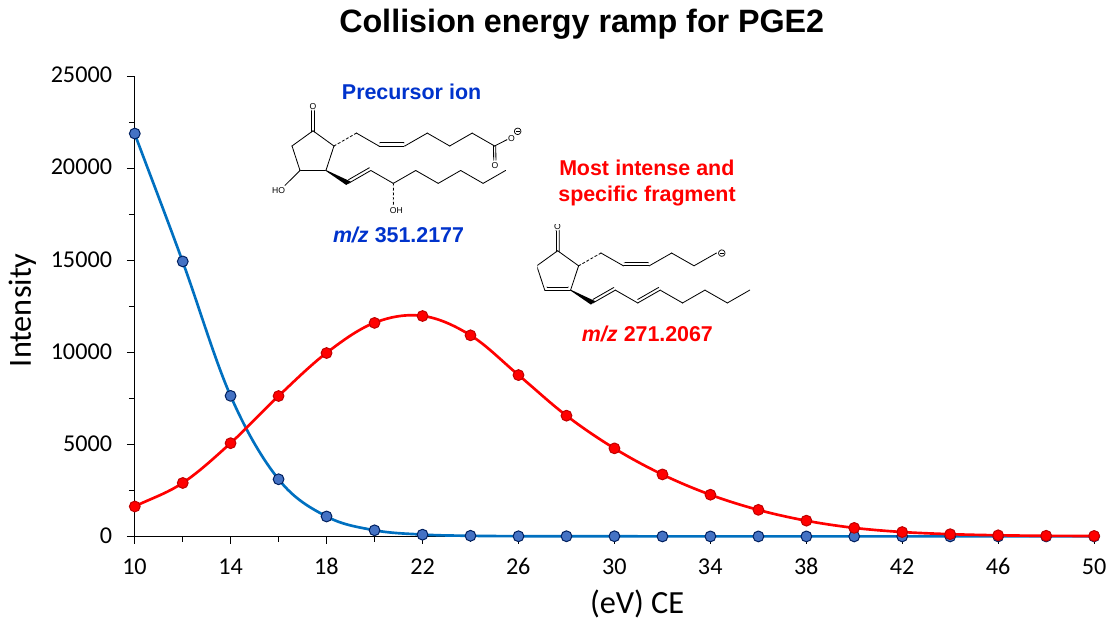
 **Figure S2. Representative collision energy ramp for PGE_2_.**
